# Single and Multiple Buds Form characteristics in *Amorphophallus muelleri*

**DOI:** 10.1101/2025.04.14.648843

**Authors:** YaXin Liu, ZeMei Li, JinDi He, WenHan Li, YuQi Xie, SiYi Ge, RuiJia Wang, XueWei Wu

## Abstract

This study compared the physiological characteristics and transcriptomic profiles of single-budded and multi-budded corms, as well as developmental stages of foliar bulbils in *Amorphophallus muelleri*, to reveal regulatory mechanisms underlying bud development. Hormonal analysis showed that the IAA/CTK ratio was significantly higher in single-budded corms (1.9060 vs 1.5753), while multi-budded corms exhibited higher CTK (0.2766 μg/g), BR (0.8692 p mol/g), and SLs (0.5090 ng/g) levels. During foliar bulbils development, ABA content peaked at 41.54 ng/g in the G2 stage, and sucrose reached maximum levels (6.86 mg/g) in the G1 stage. Transcriptomic sequencing identified 742 differentially expressed transcription factors, with over 80% of *CRE1* and *AHP* genes in CTK signaling pathways significantly upregulated. KEGG pathway analysis highlighted starch/sucrose metabolism (215 DEGs) and plant hormone signal transduction (243 DEGs) as key regulatory networks. The study demonstrated that CTK signaling activation and carbohydrate metabolic reprogramming promoted multi-bud formation, while foliar bulbils development displayed stage-specific gene expression patterns, providing theoretical insights for improving konjac propagation coefficient.

## 1. Introduction

Konjac is a perennial herbaceous species of *Amorphophallus* in Araceae (Zhang et al., 2022; Liu et al., 2019a). The underground corms of konjac are rich in konjac mannan (Behera et al., 2017; Huang et al., 2016; Guo et al., 2022). Glucomannan is a polymeric water-soluble polysaccharide (Smirnova et al., 2002) with good biodegradability, stability, and film-forming properties, which can be used to improve the functional properties of proteins (Zhang et al., 2014). In addition, it is now widely used in medicine, aviation, food, and chemical industries (Kobayashi et al., 2002; Liu et al., 2019b).

The most widely cultivated species of konjac are *A. konjac*, *A. albus*, and *A. bulbifer*. Among them, *A. konjac* and *A. albus* have the largest cultivation area in China, with a total area of 1,300 km^2^. They have excellent quality, but these two types of konjac have weak resistance to konjac soft rot disease. (Wei et al., 2020). *A. bulbifer* is a recently developed and cultivated konjac species that has been cultivated. The cultivation area is almost 400 km^2^. *A. erubescens*, *A. bulbifer*, *A. yuloensis,* and *A. muelleri* are some species of konjac that can form foliar bulbils on the petiole and lobed leaf bifurcation (Zhang et al., 2009; Hetterscheid W, 1996). The wild species of bulbil konjac is distributed in the tropical rainforest and can survive in hot and humid environments. Compared to *A. albus* and *A. konjac*, *A.bulbil* has a higher resistance to soft rot disease and a shorter growth cycle. The propagation coefficient is high, making it suitable for large-scale planting popularization (Wang et al., 2022; Su et al., 2022).

Due to the constant increase in demand for konjac mannan in the international market, konjac is in short supply throughout year (Wang et al., 2023). However, the growth cycle of konjac underground corms is long, and it takes 2 - 3 years to grow from small corms to commercial corms, increasing the cost of konjac production. *A. konjac* and *A. albus* are typically single-leaved plants. Each corm produces only one compound leaf per growing season, whereas bulbil konjac produces several leaves per growing season, and the leaves can grow foliar bulbils at their forks. Plants propagated by foliar bulbils and seeds are mostly grow on the same corm, whereas plants propagated by underground corms mostly grow on the different corm. (Zhang et al., 2009).

During the cultivation of konjac corms, a single corm can typically only produce one shoot sheath and then one compound leaf with pinnate leaflets. However, there is a phenomenon in which a corm can produce multiple sprouts, which then develop into multiple leaves and grow continuously. In addition, it can increase the leaf area and growth period of konjac in a single season, as well as facilitate the rapid expansion of the corms (Zhou et al., 2010). However, the occurrence of multiple leaves in konjac is unpredictable. The incidence of multiple leaves in corms of different species and ages appears to be different. Consequently, it is important to investigate the occurrence of multiple buds in konjac.

Numerous species within the Araceae family exhibit the development of multiple leaves during their growth process. The cultivation of konjac typically results in the production of a single compound leaf annually due to strong apical dominance (Zhou et al., 2010). In instances where the terminal bud of konjac is damaged, the lateral buds on the corm will continue to grow in place of the terminal bud. Moreover, if both the terminal and lateral buds are damaged, adventitious buds will gradually sprout and develop (Cai et al., 2020). Removal of the terminal buds in both *A. konjac* and *A. albus* has been observed to increase the number of petioles, with *A. albus* showing an increase to 1.5 petioles per corm and *A. konjac* to 2 petioles per corm (Qin, 2004). The production of only one compound leaf per growth cycle in konjac is regulated by endogenous hormones. Specifically, when the ratio of cytokinin (CTK) to growth hormone reaches a certain threshold, it can inhibit apical dominance and lead to the occurrence of multiple leaves in konjac. Experimental findings have indicated that 6-BA is beneficial in stimulating the growth of lateral buds and leaf formation in konjac (Chen et al., 2005).

Nutrients and hormones are crucial factors in the regulation of apical dominance, with auxin (IAA) being synthesized in meristematic tissues or young plant organs and then transported through polar transport systems to achieve differential distribution and influence morphogenesis (Kebrom, 2017; Ruegger et al., 1997; Friml, 2003). Plant growth hormones, such as CTK and strigolactone (SLs), have been identified as regulators of axillary bud growth, with SLs also playing a role in seed germination and the development of leaves, lateral buds, and roots (Gomez-Roldan et al., 2008). Recent research has shown that brassinolide (BR) directly influences the expression of *BZC1* through *BZR1* to mediate tomato bud growth, as well as regulating other hormones and sugars to impact axillary bud development. This highlights the potential of BR signaling in axillary buds as a target for plant morphogenesis, affecting various aspects of plant life including canopy structure, cell growth, and division (Planas-Riverola, 2019).

According to the concept of apical dominance, the growth of stems induced by IAA indirectly inhibits the growth of terminal buds. Recent research has indicated that the dormancy of terminal buds in response to internal and external factors is linked to the sugar levels present in the terminal bud. This discovery supports the idea that stem growth stimulated by growth diverts sugar away from the terminal bud, consequently impeding its growth (Kebrom et al., 2017). Some scholars suggest that the tip of the stem of a plant sustains growth by having preferential access to nutrients, which flow towards the area with the highest concentration of growth hormone (Irwin et al., 1996). Investigations on peas have demonstrated that sugar is essential and adequate for releasing apical dominance, which primarily involves restricting sugar access to axillary buds. As terminal buds in growth compete with axillary buds for sugar and hinder the growth of axillary buds through signaling, it has been proposed that sugar levels regulate the hormonal system that governs apical dominance (Schneider et al., 2019). Experiments on potato tubers have shown that subjecting them to low temperatures (4 °C) versus high temperatures (33 °C) increases tuber sugar content while promoting more stem growth. This suggests that high sucrose levels in thin tissues may trigger the release of apical dominance, potentially through sugar signaling induction. However, the triggering factor may not be limited to sucrose and could also involve hexose (Salam et al., 2017).

To enhance the reproductive capacity of *A. muelleri*, our study delved deeply into the developmental morphological characteristics, transcriptome data, and physiological indices of foliar bulbils at different stages. Meanwhile, the growth and developmental differences between single-bud corms and multi-bud corms of *A. muelleri* were analyzed. This research is supported by transcriptome data and physiological markers. The objective is to investigate the developmental process of foliar bulbils and the changes in the number of buds during the germination process of foliar bulbils, with the aim of increasing the yield of *A. muelleri* within a single growing season, thereby improving the propagation coefficient of *A. muelleri* and providing a theoretical basis for guiding the future commercial cultivation practice of *A. muelleri*.

## 2. Materials and methods

### 2.1. Plant materials

Foliar bulbils of *A. muelleri* were collected at Lincang Dian Sheng Agricultural and Forestry Development Co., Ltd. The foliar bulbils weighing 6 - 30 g were chosen and stored at a constant temperature of 26 °C in the laboratory for one month. When the buds of the foliar bulbils started to germinate, they were placed in a thermostat at 33 °C for germination treatment. Subsequently, they were planted in diameter of 30 cm flowerpots with peat soil, 5 corms per pot, followed by conventional cultivation management. During the germination period of *A. muelleri* corms, single buds and multiple buds were selected as materials. In addition, during the growth period of foliar bulbils, foliar bulbils at different developmental stages were selected as materials. After being cut out the buds and foliar bulbils, one part was preserved in FAA fixative for subsequent paraffin sectioning experiments. The remaining part was sliced, wrapped in tin foil, quickly frozen in liquid nitrogen and stored at -80 °C for transcriptome sequencing, hormone, and sugar content.

### 2.2 Histological Analysis

First, 1 cm samples were taken from the single-bud, multi-bud, and foliar bulbils, and then fixed in FAA fixation buffer (formaldehyde: glacial acetic acid: 70% ethanol, 1:1:18) for 48 h at 4 °C. The fixed samples were dehydrated in a graded series of ethanol (70%, 85%, 95%, and 100%), followed by a xylene/ethanol series (xylene: ethanol 1:2, 1:1, and 2:1 and 100% xylene). Xylene was gradually replaced with paraffin (melting point of 58 °C - 60 °C) at 60 °C for three days. Sections (10 µm thick) were obtained using a rotary microtome and were double-stained with 5% (w/v) Safranine T and 0.5%(w/v) Fast Green FCF.

### 2.3 Determination of hormone content

The content of IAA, CTK, BR, and SLs was determined in the corms of konjac using a double antibody sandwich assay with an enzyme-linked immunoassay kit supplied by Normin Koda (Wuhan) Biotechnology Co.

According to the kit’s instructions, the determination procedure was carried out, and 0.1∼0.2 g of the sample was weighed and recorded. A quantity of PBS, pH 7.4, was added, and the specimen was homogenized by hand or with a homogenizer. Approximately 20 minutes of centrifugation (2000-3000 rpm). The supernatant was carefully collected. Standard tubes and sample tubes were prepared, and 50 μL of standards with varying concentrations were added to each standard tube. The control were prepared (no samples or enzyme reagents were added; the remaining steps were identical), and the sample tubes were to be evaluated. Add 40 μL of sample diluent to the wells of the sample to be tested on the enzyme labeling plate, followed by 10 μL of the sample to be tested (resulting in a 5-fold dilution). The sample was added to the bottom of the wells of the plate, taking care not to touch the tubes, and gently mixed. The plate was sealed with sealing film and incubated at 37 °C for 30 min. The 20-times concentrated cleaning solution was diluted with 20 times its volume of distilled water and is now ready for use. The sealing film was removed with care, and the liquid was discarded. The wells were filled with washing solution, left for 30 s, and then discarded. The preceding steps were repeated five times and then dried. Subsequently, 50 μL of enzyme reagent was added to each well, excluding the blank wells. Seal the plate with a film and incubate at 37 °C for thirty minutes. The sealing film was removed carefully; the liquid was discarded, the wells were filled with washing solution, left for 30 s, and then discarded. The preceding steps were repeated five times before being patted dry. The color was developed at 37 °C for 15 minutes after adding 50 μL of Colorant A and 50 μL of Colorant B to each well, gently shaken, and mixed thoroughly. Then, 50 μL of termination solution was added to each well to terminate the reaction (at this point, the blue color will turn to yellow), the absorbance (OD value) of each well was measured sequentially at 450 nm, and a blank well was used to set the absorbance to zero.

### 2.4 Determination of sugar content

The fructose and sucrose content in samples was determined using the fructose and sucrose determination kit supplied with Nominco (Wuhan) Biotechnology Co. The determination procedure was carried out according to the kit instructions. The experiment involved weighing 0.1∼0.2 g of sample, recording the weight, adding the extraction solution, performing a water bath, centrifugation, decolorization, and then separating the extract supernatant for analysis.

### 2.5 RNA-seq and data analysis

Raw image data obtained from sequencing were converted into raw reads by base calling, and the reads containing adapter, N ratio greater than 5%, and low quality (the number of bases with quality value *Q* ≤ 10 comprised more than 20% of the entire read) were removed to obtain clean reads. De-novo assembly of clean reads was carried out using Trinity software. The reads with a specified overlap length were concatenated into a longer fragment Contig, which was then recompared with the clean reads. The transcript to which the Contig belonged and its distribution in the transcript were determined by the paired-end reads. The Trinity software was able to concatenate these contigs to obtain the two ends of the non-expandable unigene.

The unigene sequences were aligned to the protein databases NCBI non-redundant database (Nr), Swiss-Prot, the Kyoto Encyclopedia of genes and genomes database (KEGG), and the Cluster of orthologous groups of proteins (COG) (e < 10^-5^) to obtain the protein with the highest sequence similarity to a given unigene. Consequently, the protein functional annotation information for that unigene was obtained.

Based on the Nr annotation information, the GO annotation information of the unigene was obtained using Blast2 GO software, and the GO function classification statistics of all unigenes were done using WEGO software. The metabolic pathway annotations of unigene were obtained by aligning unigene sequences to KEGG database via Blastx.

After calculating the expression of unigenes using the FPKM method, differential expression analysis was performed using DESeq to identify unigenes with a padj (corrected *P*-value) 0.05 as differential genes. GO enrichment analysis with GOseq and Pathway enrichment analysis with KOBAS (2.0) were applied to differentially expressed genes identified by screening *P* < 0.05 indicated a significant enrichment.

### 2.6 Statistical Analysis

The data for fructose, sucrose, and endogenous hormones contents and the relative expression analyses of genes were presented as the means with standard errors from three biological replicates. All statistical analyses were performed using IBE SPSS Statistics 26, and statistical significance was set at *P* < 0.05. Non-overlapping letters (a-d) indicated significant differences between the comparisons based on the ANOVA analysis and Duncan’s multiple range test. The GraphPad Prism 5.0 software was used for graphics.

## 3. Results

We can distinguished single-bud from multi-bud corm by the number of its buds (Fig. 1 A-D), then we cut the bud of the corm for paraffin section (Fig. 1 E-K). And we divided the developmental process of the foliar bulbils into four periods based on its different morphological forms: the foliar bulbils pre-development period (G1), the foliar bulbils initiation period (G2), the foliar bulbils expansion period (G3), and the foliar bulbils maturation period (G4) (Fig. 1 P-S)., then we also prepared paraffin sections of the foliar bulbils at these four periods (Fig. 1 L-O).

**Fig. 1.**
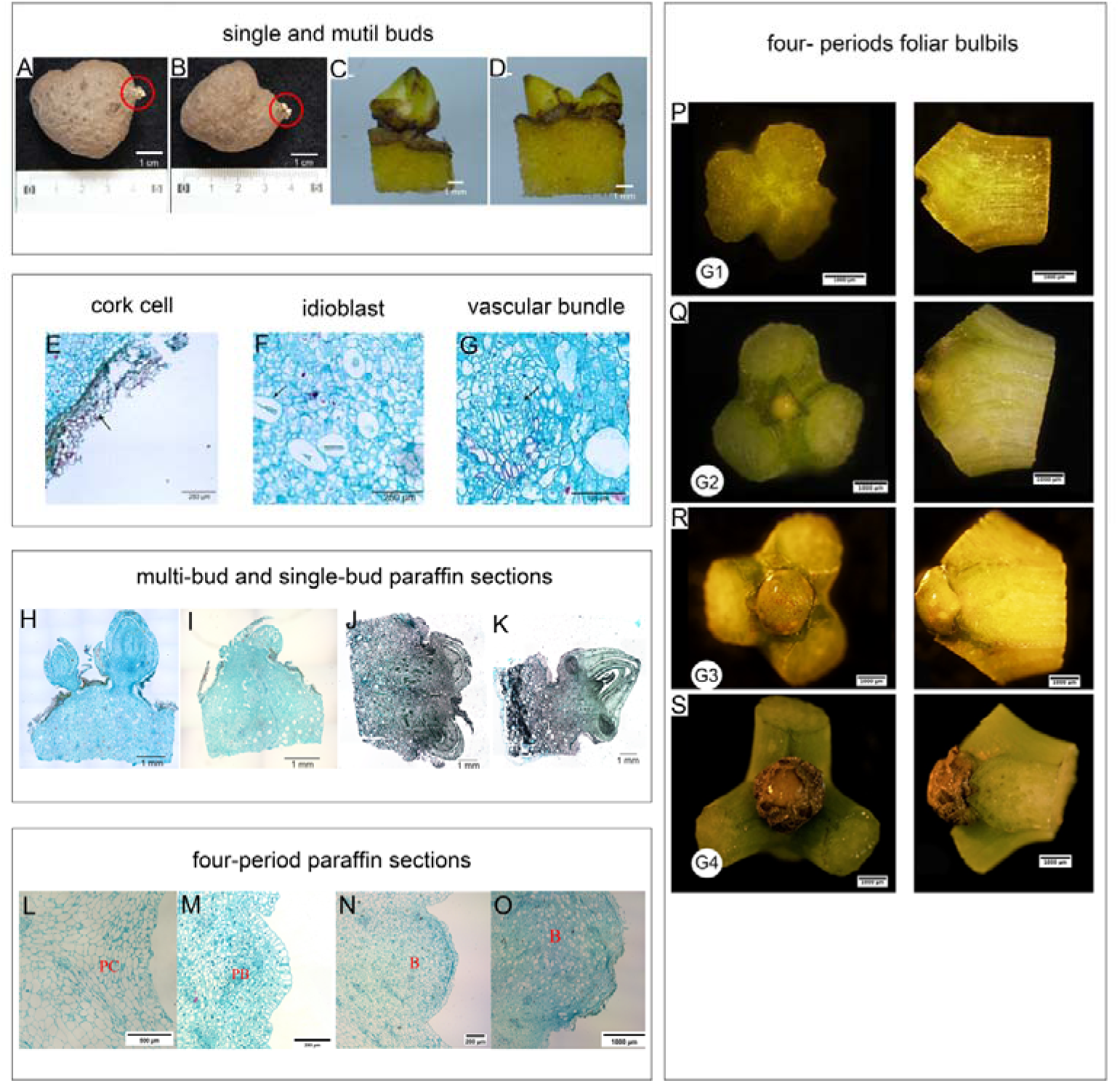
Paraffin section sampling site and paraffin section view. Notes: **A, C**: material of multi-bud;**B, D**: material of single-bud; **E**: Cork cell; **F**: Idioblast; **G**: Vascular bundle; **H-K**: The microscopic structure of the longitudinal section at the junction of bud and bulbil.(**I, K**: Material of single-bud; **H, J**: Material of multi-bud); **P-S**:The G1, G2, G3, and G4 periods and their cross-sectional views of foliar bulbils; **L-O**: The microscopic structure of the longitudinal section at the junction of foliar bulbils. (**PC**: Parenchyma cell**; PB**: Bulbil primordium; **B**: Bulbil).

### 3.1 Morphological observation

The longitudinal sectioning of the corms-bud junction revealed that its composition included the cortex, phellogen, thin-walled cells, vascular bundles, phloem tissue, and buds. Meanwhile, the longitudinal sectioning of the foliar bulbils at the junction of the compound leaf petiole revealed its composition to include the cork layer, thin-walled cells, and vascular bundles.

The foliar bulbils of *A muelleri*’s grows at the fork of the petiole of compound leaves and experiences long-term exposure to sunlight. The outer cortex displays an uneven and coarse texture, appearing brown in color. The cork layer cells are predominantly rectangular in shape (Fig. 1 E, 1 S). During the foliar bulbils development and post-maturation sprouting stages, the interior of the corm is predominantly composed of parenchyma tissue and vascular tissue(Fig. 1 L). The parenchyma cells store nutrients essential for supporting the initial growth and development of the plant. Notably, the volume of idioblasts containing needle crystals exceeds that of the surrounding parenchyma cells, with the idioblast cytoplasm exhibiting minimal staining (Fig. 1 F). Additionally, during the development of the bulbils at the base of the petiole, the vascular bundles are primarily located within the petiole. From the G1 to G4 periods of bulbil development, organic matter begins to accumulate around the vascular bundles. In the G4 period, both transverse and longitudinal vascular bundles maintain the nutrient transport system, ensuring the development and maturation of the bulbils. (Fig. 1 G, M, N).

By examining the paraffin sections of single and multiple buds, we observed a large amount of vascular tissue beneath the bud and the bulbous leaf, with both transverse and longitudinal orientations (Fig. 1 K). The outer region of the bud contains young leaves, while the inner part of these leaves exhibits numerous large cells located near the periphery, with a volume several times greater than the surrounding cells. Conversely, the inner portion consists of relatively small cell clusters. Beneath the young leaves, meristem cells with large nuclei, tightly packed arrangement, and diminutive cell size were identified, indicating active cell division. The transition from the lower section of the young leaves towards the corm’s interior demonstrates a pattern of diminishing nuclear visibility and increasing cell size (Fig. 1 I). Through paraffin section view of foliar bulbils observations, it was found that from the G3 period, the epidermal cells began to further widen and thicken(Fig. 1 N). The meristematic cell clusters beneath the epidermis continued to divide and proliferate, leading to an increase in the number of internal cells, causing the base of the petiole to expand and protrude, forming the bulbil structure (Fig.1 O). By comparing cross-sections from the G2 and G3 periods, it is evident that starting from the G3 period, the organic matter content in the cells surrounding the vascular bundles significantly increased, indicating that organic matter is being transported to the developing bulbil region(Fig. 1 M, 1 N).

Numerous disorganized vascular bundles were also observed below the multi-bud material (Fig. 1 J). The observation reveals that the central two layers of nascent leaves in the larger bud are enveloped by cells characterized by a diminutive size and a prominent nucleus. The cells situated at the lower periphery of the cellular stratum exhibit a resemblance to those in a solitary bud, being several times larger than their neighboring cells, while those positioned more internally are comparatively smaller in size. Concurrently, in the lower region of the larger bud, three meristematic cells displaying a diminutive cellular volume, distinct nucleolus, close proximity, and robust division activity were identified. The juxtaposed left and right clusters exhibit a superimposed configuration akin to that of a bud, suggesting a potential identity as an axillary bud (Fig. 1 H).

### 3.2 Determination of Physiological Indicators

#### 3.2.1 Determination of endogenous hormone content

Hormone determination was carried out on single - bud, multi - bud, and foliar bulbils at different developmental stages. There was no significant difference in the IAA content between single - bud konjac corms and multi - bud konjac corms (*P* > 0.05), which were 0.4299 μg/g and 0.4349 μg/g, respectively. However, the CTK content in multi - bud corms was significantly higher than that in single - bud corms (*P* < 0.05), being 0.2766 μg/g and 0.2556 μg/g, respectively. In addition, by calculating the ratio of IAA to CTK, it was found that the IAA/CTK ratio in single - bud corms was significantly higher than that in multi - bud corms (*P* < 0.05), with values of 1.9060 and 1.5753, respectively.

The overall trend of ABA was to increase first and then decrease slowly. The ABA content in the G1 period was only 11.29 ng/g, but in the G2 period, it increased nearly four - fold to reach the maximum value of 41.54 ng/g. In the G3 period, it remained at a relatively high level of 39.75 ng/g. The ABA content in the G4 period was significantly lower than that in the G2 period (*P* < 0.05), being 27.40 ng/g.The IAA and CTK contents and their ratios of single-bud and multi-bud corms are shown in (Fig. 2 A-C). Fig. 2 D-E displays the contents of BR and SLs in single and multi-bud corms. The BR content in single-bud corms was significantly higher than that in multi-bud corms (*P <* 0.05), with mean values of 1.3111 p mol/g and 0.8692 p mol/g, respectively. The SLs content in single-bud corms was significantly higher than that in multi-bud corms (*P <* 0.05). The mean values were 1.0862 ng/g and 0.5090 ng/g, respectively.

**Fig. 2.**
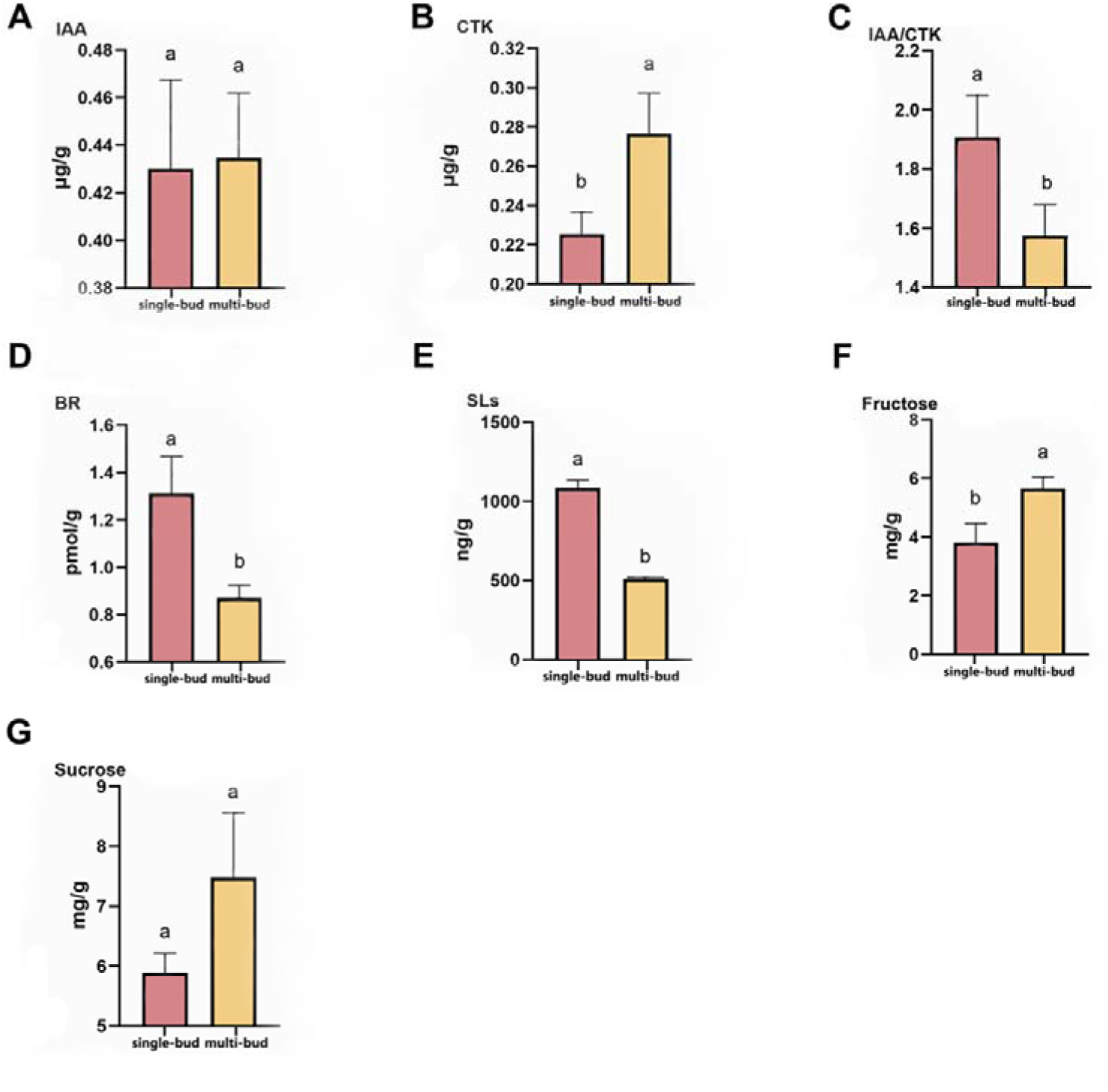
Hormone, fructose and sucrose content in single and multi-bud corms. Notes: **A**: IAA content; **B**: CTK content; **C**: IAA to CTK ratio; **D**: BR content; **E**: SLs content; **F**: fructose content; **G**: sucrose content; different letters represent significant differences between samples (*P <* 0.05).

#### 3.2.2 Determination of sugar content

The contents of sucrose and fructose were determined in single buds, multiple buds, and foliar bulbils at different developmental stages. The fructose and sucrose contents in single - bud and multiple - bud corms are shown in (Fig. 2 F - G). There was no significant difference in sucrose content between single - bud and multiple - bud corms (*P* > 0.05), with average values of 5.8893 mg/g and 7.4809 mg/g, respectively. In contrast, the fructose content was different. The fructose content in multiple - bud corms was significantly higher than that in single - bud corms (*P* < 0.05), with values of 5.6623 mg/g and 3.8093 mg/g, respectively. For the determination of foliar bulbils at four developmental stages, the sucrose content is shown in (Fig. 3 K). The results indicated that the sucrose content was the highest, 6.86 mg/g, at the G1 period, and decreased by nearly 30% to 4.80 mg/g at the G2 period. During the G3 period and the G4 period, the sucrose content increased slowly. There was no significant difference in sucrose content among the four stages (*P* > 0.05). This suggests that a high concentration of sucrose may play a crucial role in the induction of bulbils, and sucrose regulates the formation and development of bulbils by acting as a signaling molecule.

**Fig. 3.**
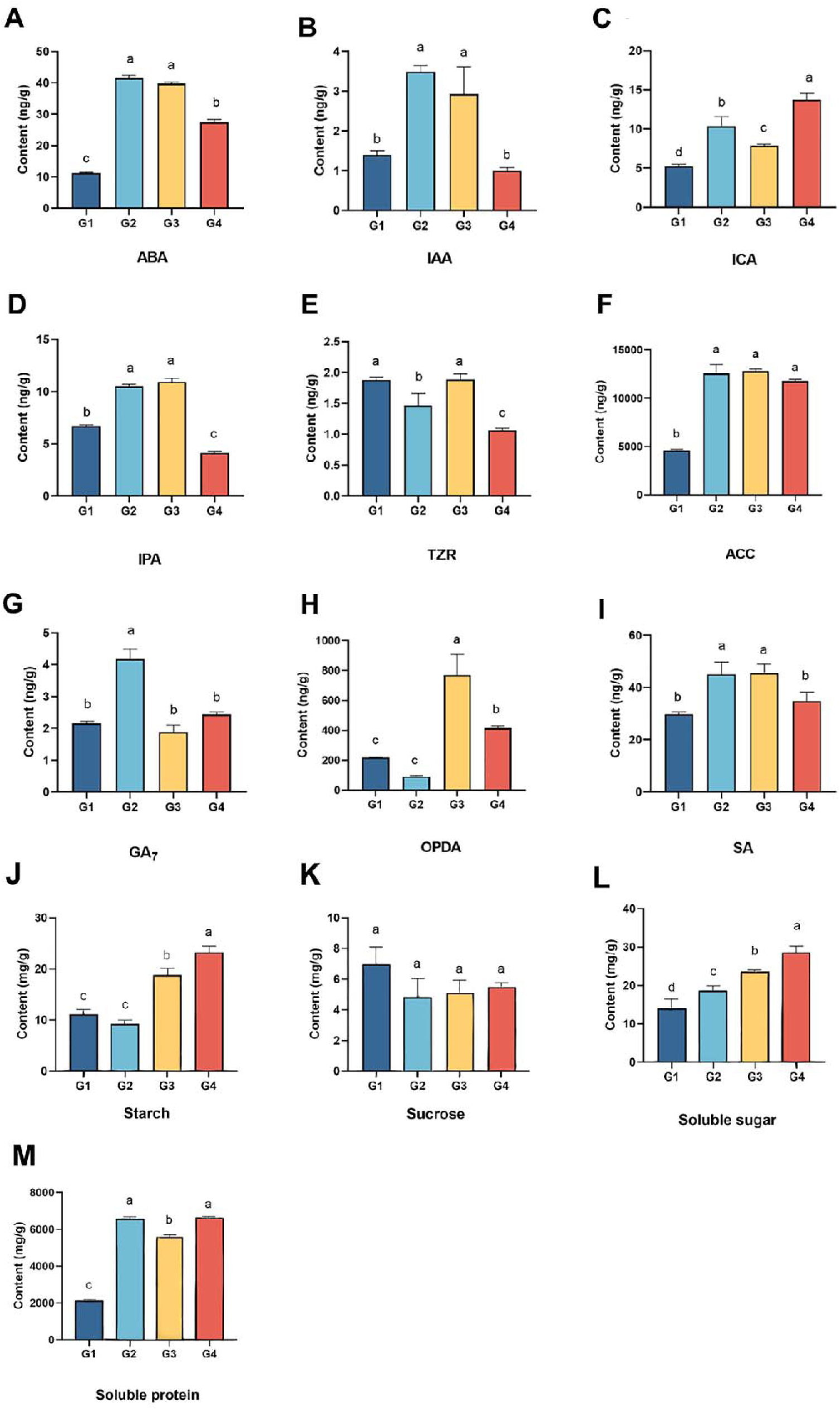
Endogenous, fructose and sucrose hormone content in different developmental periods of bulbil Note: **A**: ABA content; **B**: IAA content; **C**: ICA content; **D**: IPA content; **E**: TZR content; **F**: ACC content; **G**: GA content; **H**: OPDA content; **I**: SA content; **J**: starch content; **K**: sucrose content; **L**: soluble sugar content; **M**: soluble protein content. Different letters indicate significant differences among samples (*P* < 0.05).

### 3.3 Transcriptome analysis of bud corm and foliar bulbils

#### 3.3.1 RNA-seq data profile

We conducted data filtering and quality control on the raw data of RNA sequencing for buds in different quantities (Group 1) and leaf surface spheres at four different developmental stages (Group 2), respectively. The results showed that the number of clean reads in each library ranged from 3.9×10L to 5.1×10L. The sequencing error rates of the two transcriptomes were both less than 0.03%. The Q20 and Q30 values were higher than 97% and 91.54%, respectively, indicating that the sequencing quality of the two transcriptomes was high and suitable for further analysis (Table 1).

**Table 1.** Statistics of sequencing output.

| Transcriptome | Sample | Reads NO. | Bases (G) | N(%) | Q20(%) | Q30(%) |
| --- | --- | --- | --- | --- | --- | --- |
| Bud | single-bud 1 | 44,108,588 | 6.62 | 0.03 | 97.59 | 92.92 |
|  | single-bud 2 | 46,798,174 | 7.02 | 0.03 | 97.56 | 92.47 |
|  | single-bud 3 | 40,571,160 | 6.09 | 0.03 | 97.77 | 93.36 |
|  | multi-bud 1 | 47,710,190 | 7.16 | 0.03 | 97.24 | 91.54 |
|  | multi-bud 2 | 39,045,196 | 5.86 | 0.03 | 98.01 | 94.07 |
|  | multi-bud 3 | 47,438,528 | 7.12 | 0.02 | 98.25 | 94.54 |
| Foliar bulbils | G1-1 | 46,363,888 | 7.00 | 0.00 | 97.49 | 93.46 |
|  | G1-2 | 44,641,622 | 6.74 | 0.00 | 97.18 | 92.83 |
|  | G1-3 | 50,477,016 | 7.62 | 0.00 | 97.34 | 93.21 |
|  | G2-1 | 41,265,928 | 6.23 | 0.00 | 97.72 | 93.74 |
|  | G2-2 | 46,311,460 | 6.99 | 0.00 | 97.71 | 93.82 |
|  | G2-3 | 49,162,872 | 7.42 | 0.00 | 97.39 | 92.94 |
|  | G3-1 | 48,214,766 | 7.28 | 0.00 | 97.22 | 92.91 |
|  | G3-2 | 48,985,970 | 7.40 | 0.00 | 97.29 | 93.04 |
|  | G3-3 | 47,795,118 | 7.22 | 0.00 | 97.23 | 92.95 |
|  | G4-1 | 49,888,026 | 7.53 | 0.00 | 97.33 | 93.16 |
|  | G4-2 | 50,449,590 | 7.62 | 0.00 | 97.25 | 92.94 |
|  | G4-3 | 44,559,464 | 6.73 | 0.00 | 97.43 | 93.29 |

The clean reads were spliced using Trinity software to obtain the reference sequences for subsequent analysis. The results yielded 286,934 transcripts with an average length of 815 bp, N50 length of 1,544 bp, and N90 length of 283 bp. The transcripts were hierarchically clustered using Corset software to obtain Clusters. In contrast, clusters with the longest sequences were classified as unigene, yielding unigene 152,464 with an average length of 1,082 bp, N50 length of 1,768 bp, and N90 length of 444 bp; For the transcriptomes of foliar bulbils in different periods, a *de novo* assembly strategy was adopted for subsequent analysis. The clean reads were assembled. The total number of transcript sequences was 356,101, with an N50 of 1,912 bp. After assembling all the transcripts, 145,629 non - redundant unigenes were obtained, with an N50 of 1,331 bp (Table 2).

**Table 2.** Statistics of assembly results.

| Type | Sequence Number | N50(bp) | N90(bp) | Total length (bp) |
| --- | --- | --- | --- | --- |
| Transcript -bud | 286,934 | 1,544 | 283 | 233,957,467 |
| Unigene -bud | 152,464 | 1,768 | 444 | 164,920,354 |
| Transcript - foliar bulbils | 356,101 | 1,912 | 493 | 433,806,502 |
| Unigene - foliar bulbils | 145,629 | 1,331 | 388 | 133,080,318 |

Unigene sequences were functionally annotated to obtain the functional information of the genes. Annotation was performed using KEGG, Nr, SwissProt, TrEMBL, KOG, GO, and Pfam (Table 3). The TrEMBL database received the most annotations, 54.35%, followed by the Nr and GO databases, 54.35%, accounting for 54.16% and 45.49%, respectively. In addition, the SwissProt had the least annotated information, accounting for only 34.02%, while 93,454 unigene were annotated to at least one of the aforementioned databases, accounting for 61.3%. In the transcriptome of foliar bulbils, an exhaustive alignment analysis was carried out on 145,629 unigenes. Through sequential alignments with seven public databases, precise functional annotations were performed. The results are presented in Table 3: 47,897 (32.89%) unigenes were mapped to NR, 25,443 (17.47%) unigenes to GO, 18,163 (12.47%) unigenes to KEGG, 20,834 (14.31%) unigenes to Pfam, 50,380 (34.59%) unigenes to eggNOG, and 32,173 (22.09%) unigenes to Swissprot.

**Table 3.** Annotation statistics.

| Database | Number - bud | Percentage- bud (%) | Number- foliar bulbils | Percentage- bud (%) |
| --- | --- | --- | --- | --- |
| KEGG | 63,433 | 41.61 | 18,163 | 12.47 |
| NR | 82,581 | 54.16 | 47,897 | 32.89 |
| SwissProt | 51,864 | 34.02 | 32,173 | 22.09 |
| TrEMBL | 82,871 | 54.35 | — | — |
| KOG | 54,038 | 35.44 | — | — |
| GO | 69,358 | 45.49 | 25,443 | 17.47 |
| Pfam | 63,950 | 41.94 | 20,834 | 14.31 |
| eggNOG | — | — | 50,380 | 34.59 |

#### 3.3.2 Sequence function annotation and function classification

Transcriptomic profiling across distinct developmental stages of bulbifer revealed conserved functional patterns in gene ontology (GO) classification and divergence in database-specific annotations. In the GO analysis of single- versus multi-bud corms (Fig. 4A), 69,358 unigenes were categorized into 47 subcategories across three domains. Molecular functions were dominated by binding (39,748) and catalytic activity (37,544), while biological processes showed enrichment in cellular (43,217) and metabolic processes (36,485). These findings were corroborated by the foliar bulb study spanning G1 - G4 stages, where 25,443 annotated unigenes exhibited similar functional proportions: binding (67%, 16,993) and catalytic activity (57%, 14,446) in molecular functions, with cellular processes (84%) and metabolic processes (72%) predominating biological processes (Fig. 4A). Cross-database comparisons revealed stage-specific annotation patterns. The corm study identified 54,038 KOG-annotated unigenes clustered into 25 groups, with general function prediction (12,317), posttranslational modification (5,979), and signal transduction (4,228) being most abundant (Fig. 4B). Conversely, foliar bulb analysis through eggNOG database annotated 50,380 unigenes into 26 functional categories, showing distinct emphasis on replication/recombination/repair (14%) and signal transduction mechanisms (8%), while 26% remained functionally uncharacterized (Fig. 4B). Notably, cellular component annotations demonstrated developmental consistency - cellular anatomical entities comprised 55,059 unigenes (93% in corms) versus 23,544 (93% in foliar bulbils), suggesting conserved structural organization across tissues. These comparative annotations highlight both stage-independent biological priorities (metabolic regulation, cellular organization) and developmental phase-specific functional adaptations in transcriptomes.

**Fig. 4.**
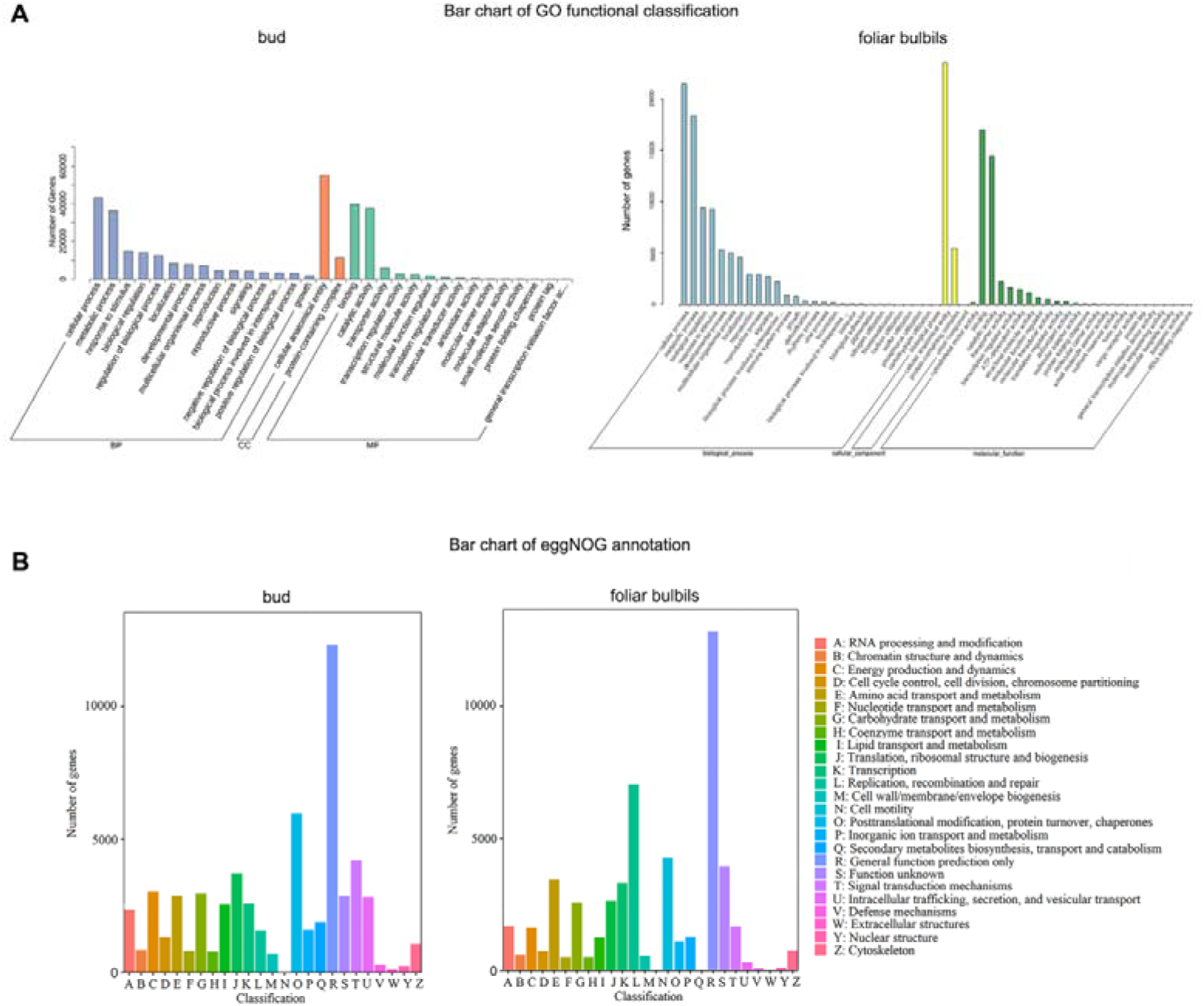
Bar chart of GO functional classification and statistics of eggNOG annotation in Transcriptome analysis of bud and foliar bulbils Notes: **A**: Bar chart of GO functional classification for bud transcriptome and foliar bulbils transcriptome; **B**: Bar chart of eggNOG annotation for bud transcriptome and foliar bulbils transcriptome.

#### 3.3.3 screening and enrichment analysis of differential genes

A total of 19,343 genes were found to be significantly different from each other, using |log2Fold Change| >= 1 and FDR < 0.05 as the threshold to screen for significantly differentially expressed genes among the samples. Compared to the single-bud samples, 10,665 genes were upregulated, while 8,678 genes were down-regulated in the multi-bud samples (Fig. 5 A-D).

**Fig. 5.**
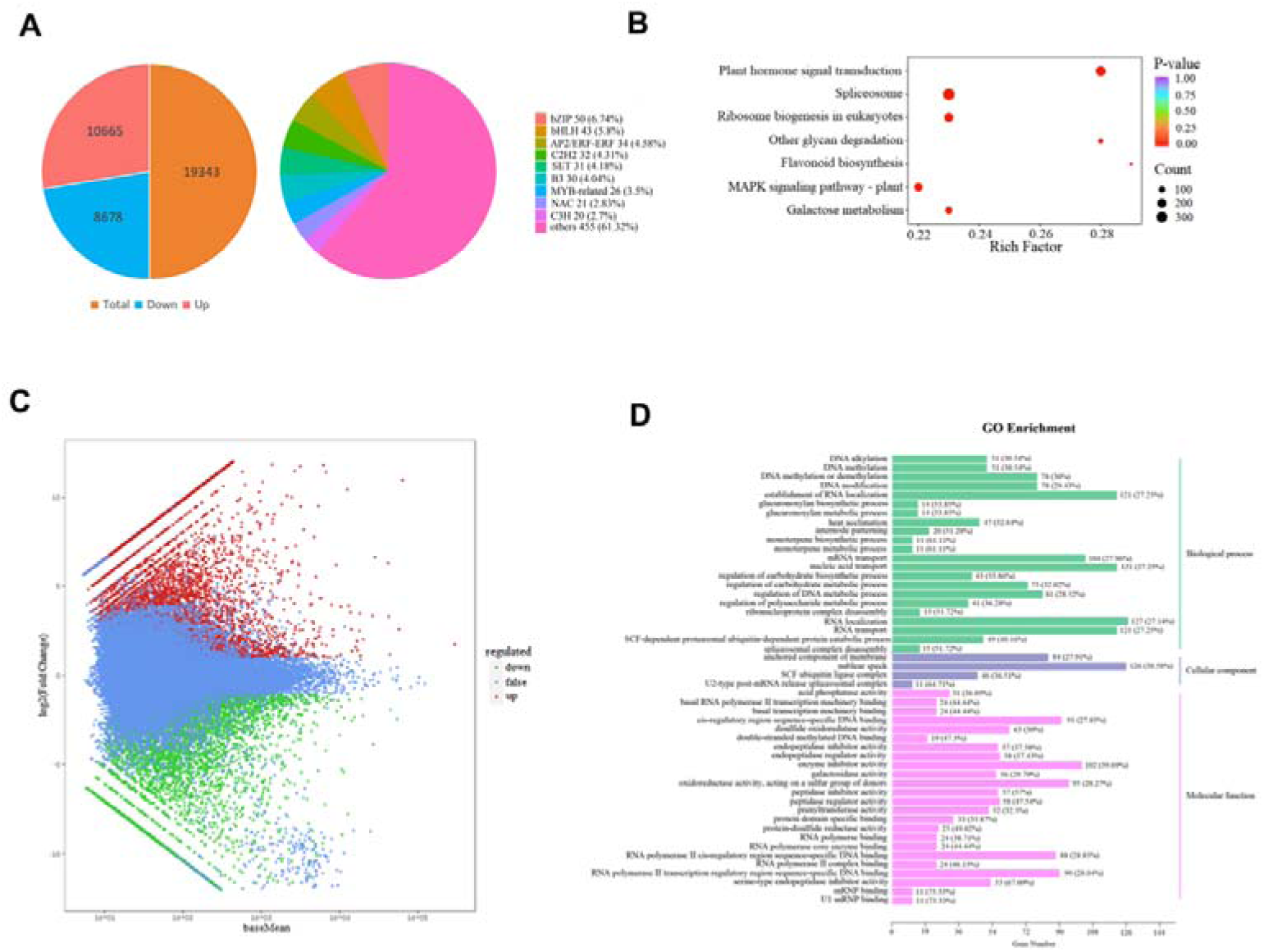
Graph of differential gene screening and enrichment analysis for bud transcriptome Notes: **A**: The number of DEGs and statistical pie charts for classification of transcription factors; **B**: Scatter plot of KEGG pathway enrichment; **C**: MA map of DEGs; **D**: Histogram of GO enrichment of DEGs.

On the DEGs of single-bud and multi-bud samples, GO enrichment analysis was performed, and the 50 GO-Term with the lowest *Q* values were selected to plot the histogram of enrichment entries (Fig. 5 D). In the biological process category, there were 65 GO-Term entries with a significance level of *Q* < 0.05, and the largest number of differentially expressed genes (DEGs) was observed for RNA localization, nucleic acid transport, RNA transport, and RNA localization establishment, totaling 127,121 DEGs. Within the cellular components category, 20 GO-terms had *Q* < 0.05, with the most notable number of DEGs identified for nuclear speck and membrane-anchored components, comprising 126 and 84 DEGs, respectively. Among the molecular functions, 56 GO-terms had *Q* < 0.05, with enzyme inhibitor activity, oxidoreductase activity, acting on a sulfur group of donors, cis-regulatory region sequence-specific DNA binding, and RNA polymerase II transcription regulatory region sequence-specific DNA binding showing the highest number of DEGs, with 102, 95, 91, and 90 entries, respectively(Fig. 5 D).

The DEGs of single-bud and multi-bud samples were analyzed for KEGG enrichment. For the enrichment scatter plot, the KEGG pathways with a *Q*-value of <0.05 were selected to plot the enrichment scatter plot (Fig. 5 B). As a result, a total of 7 KEGG pathways were significantly enriched. Among them, the spliceosome pathway contained the highest number of DEGs (differentially expressed genes) at 345, followed by plant hormone signal transduction with 243 DEGs. The remaining pathways ranked in descending order were: ribosome biogenesis in eukaryotes (196 DEGs), MAPK signaling pathway - plant (163 DEGs), galactose metabolism (120 DEGs), other glycan degradation (70 DEGs), and flavonoid biosynthesis (46 DEGs).

Based on the annotated categorical statistical data of differential transcription factors to draw pie charts (Fig. 5 A), the findings indicated that there were 742 transcription factors exhibiting significantly varied expressions between the control single-bud konjac corms and the multi-bud konjac corms. Within this group, 370 transcription factors were upregulated, while 372 were down-regulated, spanning across 79 families. Notably, the bZIP family had the highest representation with 50 members, followed by bHLH, AP2/ERF-ERF, C2H2, SET, and B3 families, with 43, 34, 32, 31, and 30 members, respectively.

In the study of the foliar bulbils transcriptome, when comparing the developmental stages of G1 and G2, a total of 5,142 differentially expressed genes were identified. Among them, 2,782 genes were upregulated, and 2,360 genes were downregulated When comparing the G2 and G3 stages, the total number of differentially expressed genes increased to 5,650, with 2,771 genes being upregulated and 2,879 genes being downregulated. In the comparison between G3 and G4 stages, the number of differentially expressed genes further increased to 7,503, with 2,743 upregulated genes and 4,760 downregulated genes. During the early stage of bulbil development, gene upregulation was the main trend, while during the late stage, gene downregulation was more prevalent. It is considered that bulbils development may be related to the up - regulated expression of certain genes(Fig. 6 A). During the entire developmental process, clust1 showed a significant upward trend in G3 and G4 compared to G1 and G2. Clust2 decreased notably from G1 to G2, then gradually increased. Clust3 increased significantly at G2 but decreased in G3 and G4. Clust4 remained stable in the first three stages and rose sharply at G4. Clust5 exhibited a trend of first increasing and then continuously decreasing across the four stages. In clust6, G2 and G3 were nearly unchanged but lower than G1, with G4 also decreasing. Clust7 increased significantly at G2, followed by continuous decreases in G3 and G4. Both clust8 and clust9 showed a marked rise at G3, with clust8’s G3 expression higher than the other three stages. Notably, in the trend plots of clust3, clust5, and clust7, G2 period was significantly higher than other stages, suggesting that genes related to bulbil formation may be highly expressed to promote bulbil germination (Fig. 6 B).

**Fig. 6.**
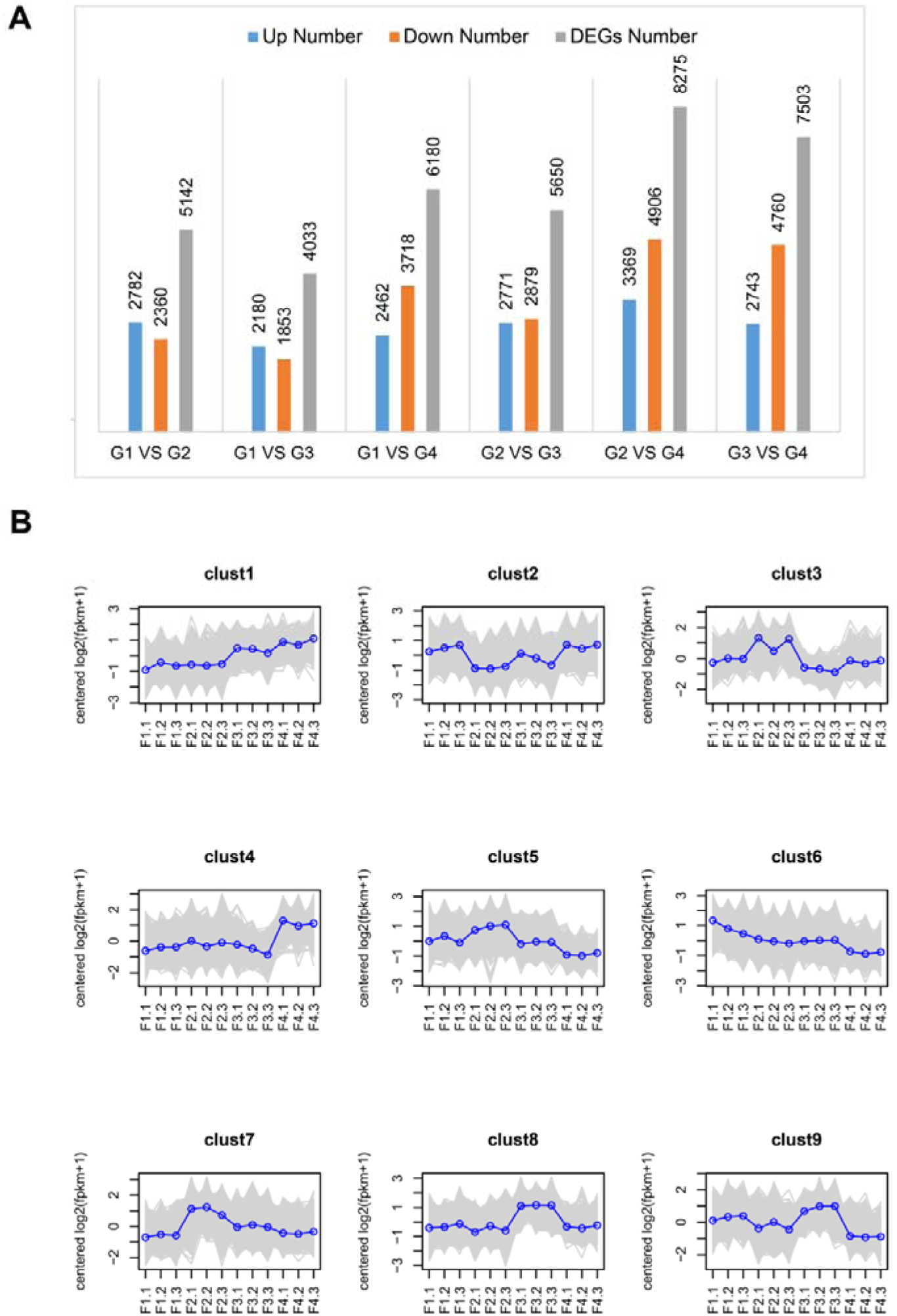
Graph of differential gene screening and enrichment analysis for foliar bulbils transcriptome Notes: **A**:Statistical results of differential expression analysis; **B**: Differential gene trend analysis.

#### 3.3.4. Analysis of differentially expressed genes associated with apical dominance

##### 3.3.4.1 Plant nutrition-related differential expression analysis

In this study, the number of differential genes annotated in KEGG for Starch and sucrose metabolism was counted. In multi-bud material compared to single-bud material, the results revealed 215 differential genes, of which 120 were upregulated. Comparatively, 95 differentially expressed genes with a *Q*-value of 1 were downregulated in this KEGG pathway, indicating that the starch and sucrose metabolic pathway was not differentially expressed between multi-bud and single-bud materials. A further search of the unigene annotation list for genes annotated as primary substances for sucrose synthesis, degradation, and transport (*SPS*, *SS*, *INV*, *SWEET*, *SUTs*) revealed the association of eight distinct genes with *SPS*. In comparison with single-bud corms, four genes in multi-bud corms showed upregulated expression, while four genes exhibited downregulated expression. Seven differential genes were associated with *SS*, with three upregulated and four downregulated. Six genes were associated with *INV*, with one upregulated and six downregulated (Fig. 7 A).

**Fig. 7.**
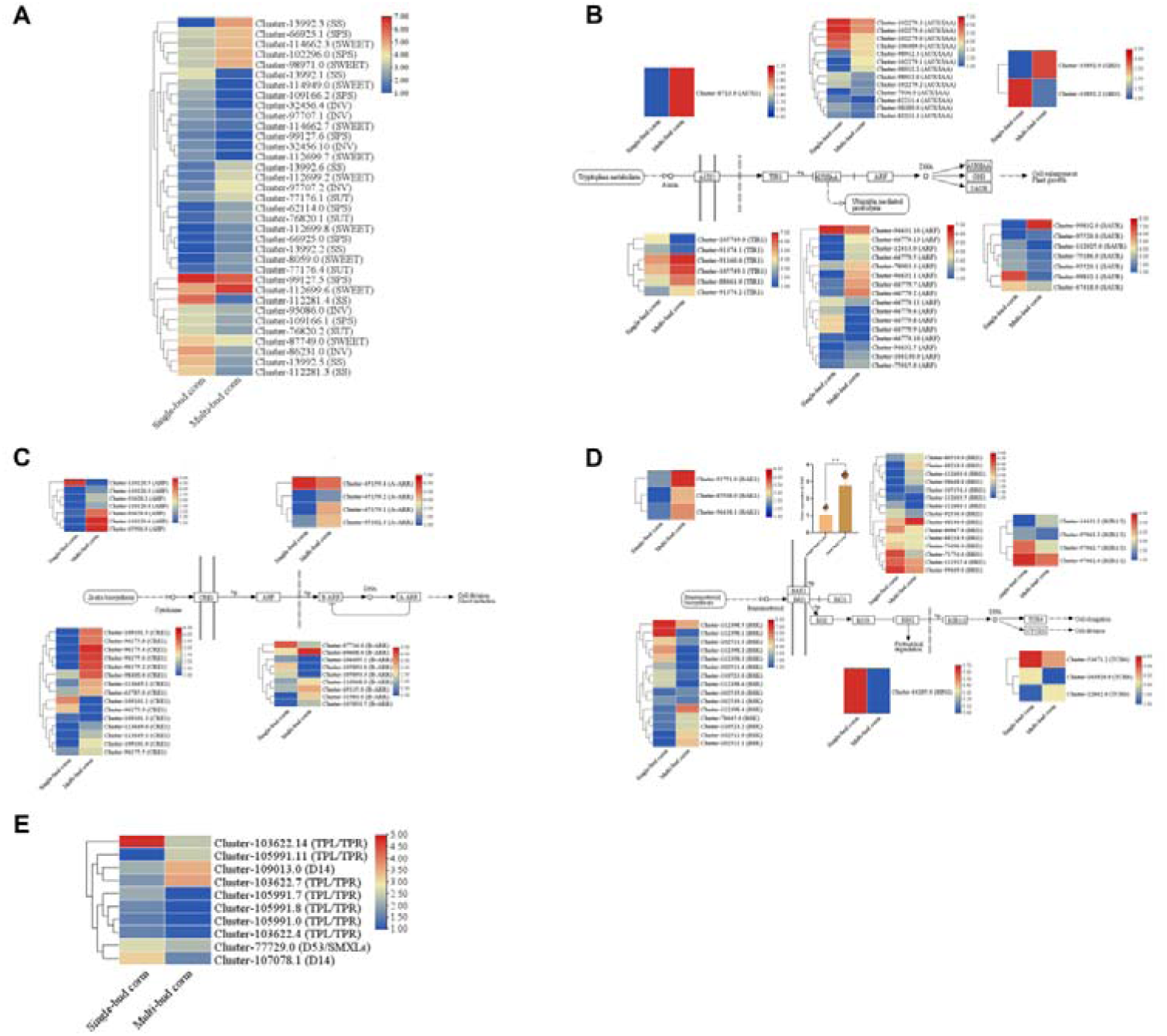
The graph of Analysis of differentially expressed genes associated with apical dominance Notes: **A**: Expression profiles of major genes involved in sucrose synthesis, degradation and transport; **B**: Expression profile of IAA signal transduction-related genes; **C**: Expression profile of CTK signal transduction related genes; **D**: Expression profile of BR signal transduction-related genes ; **E**: Expression profile of SLs signal transduction-related genes.

##### 3.3.4.2 Analysis of differential expression of plant signal transduction pathways

In this study, the number of genes involved in plant hormone signal transduction with different expression patterns was determined. Compared to single-bud material, multi-bud material contained 243 differential genes, including 133 upregulated expressed genes and 110 downregulated expressed genes. The *Q* value of this pathway was 0, making it the pathway with the lowest *Q* value among all significantly enriched KEGG pathways.

During IAA signaling, 45 differentially expressed genes were enriched, with one gene, *AUX1*, upregulated compared to single bud material, six associated with *TIR1*, of which four were upregulated and two were downregulated, 13 associated with AUX/IAA, of which four were upregulated and nine were downregulated; 16 associated with *ARF*, of which 10 were upregulated and six were downregulated; two associated with *GH3*, of which one was upregulated, and one was downregulated (Fig. 7 B).

A collective of 35 distinct genes exhibited enrichment in the context of CTK signaling. Among these, 15 genes were linked to *CRE1*, with 12 showing upregulation and three displaying downregulation in comparison to single bud material. Furthermore, seven genes were associated with *AHP*, with six showing upregulation and one showing downregulation. Additionally, nine genes were linked to *B-ARR*, with five displaying upregulation and four showing downregulation, while four genes were associated with *A-ARR*, with three showing upregulation and one showing downregulation (Fig. 7 C).

During the process of brassinosteroid signaling, a set of 41 genes exhibited enrichment. Among these, three genes linked to *BAK1* showed upregulation in comparison to single bud material, while 15 genes associated with *BRI1* displayed either upregulation or downregulation. Notably, there were 15 genes related to *BSK*, with six showing upregulation and nine displaying downregulation. Additionally, one gene was linked to the upregulated *BIN2* gene. Within the group of four genes associated with *BZR1/2*, one gene demonstrated upregulation, whereas the remaining three exhibited downregulation. Furthermore, three genes were connected to *TCH4*, with one being upregulated and two being downregulated (Fig. 7 D). Since BR have a certain regulatory effect on bud growth as demonstrated in previous studies, a q-PCR analysis was conducted on the *BAK1* gene within the BR signaling regulation pathway. The results were consistent with the transcriptome data, showing that the expression level of the *BAK1* gene in multi-bud samples was significantly higher than that in single-bud samples.

Since the unigene signaling pathway has not been fully revealed, it is not presented in the KEGG database. Therefore, in this study, we searched for genes associated with SLs signaling (*D14*, *D3/MAX2*, *D53/SMXLs*, *TPL/TPR*, and *IPA1*) in the unigene annotation list and identified the following differentially expressed genes. Compared to single-bud material, one gene related to *D53* was found to be downregulated. There were two genes related to *D14*, one upregulated and the other downregulated, and seven genes related to *TPL*/*TPR*, of which two were upregulated, and five were downregulated (Fig. 7 E).

## 4. Discussion

To improve the propagation coefficient of *A. muelleri*, this study explored the development mechanisms of foliar bulbils and multi-leaf formation in *A. muelleri*. In terms of the multi-leaf research, the vegetative organs of *A. muelleri* contain mannan and two types of storage polysaccharides. The idioblasts that produce mannan are several times larger in volume than the cells storing starch (Shi et al., 1998). In this study, it was also observed that some cells containing raphides were relatively large. When the leaves of *A. muelleri* corms emerge, the procambium differentiates from the parenchyma beneath the scales and then further differentiates into vascular tissues, which connect with the original vascular tissues in the corm to transport nutrients and water for the growth and development of plants (Liu et al., 1986). Moreover, well-developed vascular tissues were found both below the single-bud and multi-bud and inside the corm, providing a tissue basis for the regulation of bud differentiation. The synergistic effect of endogenous hormone dynamics and sugar metabolism reveals the mechanism of multi-bud formation. Endogenous hormone analysis showed that the CTK content in multi-bud *A. muelleri* was significantly higher than that in single-bud *A. muelleri* (*P* < 0.05), and the IAA/CTK ratio was significantly lower (*P* < 0.05). This was similar to the regulatory patterns in which the number of single-branch leaves of sweet potatoes increased after CTK treatment (Qiu, 2008), the CTK content in the sho and hoc mutants of *Arabidopsis thaliana* with a multi-branching phenotype was high (Xia et al., 2022), and the tillering bud growth was promoted when the IAA/CTK ratio was low (Zhang et al., 2022). The SLs content was higher in the corms of single-bud, which was consistent with the *d27* rice mutant with reduced SLs content and increased tillering (Wang, 2021) and the pea mutant with a significant reduction in SLs content accumulation and a multi-branching phenotype (Li, 2021), indicating that SLs inhibited bud differentiation. Additionally, the BR content in the multi-leaf corms at the seedling-emergence stage was extremely significantly higher than that in the single-leaf *A. muelleri* corms (*P* < 0.01), which was consistent with the regulatory role of BR in promoting tillering in rice (Jiang, 2022) and the inhibition of lateral branches in tomatoes when BR was inhibited (Gu, 2021). In the sugar metabolism of multi-leaf *A. muelleri* corms, fructose was enriched in the multi-bud corms, which was in line with the positive correlation between tiller number and fructose content in Kentucky bluegrass (Chen, 2021), suggesting that the growth of multi-buds required more fructose. The regulation of plant apical dominance is complex. After its breakdown, the interaction between sugars and hormones promotes the growth of axillary buds (Shi, 2020), and changes in hormones and corm sugars affect apical dominance. In the future, the number of buds and leaves of *A. muelleri* plants could be increased by combining hormone and sugar nutrient regulation. In the study of foliar bulbils, bulbils are mostly differentiated from parenchyma cells that regain their ability to divide. The bulbils of *Pinellia cordata*, *Pinellia ternata*, and *Dioscorea opposita* all originate from parenchyma cells (Zhu et al., 2017; Luo et al., 2014; Zhang et al., 1994). The bulbils of *A. muelleri* are formed by the differentiation of the 5th - 10th layers of parenchyma cells near the proximal end of the petiole base. In this experiment, the increase in IAA concentration promoted the initiation of bulbils and then declined after reaching a critical value, which was consistent with the conclusion of *Pinellia ternata* bulbils formation (Chang et al., 2007). IAA is related to cell growth (Roumeliotis et al., 2012) and starch synthesis (Borzenkova et al., 1998). Similar patterns were found in the formation of lily bulbs (Podwyszynska., 2006) and tulip bulbs (Zheng et al., 2012). In *A. muelleri*, the CTK, IPA and TZR reached their highest values in the G3 stage. The TZR content in the G2 stage slightly decreased due to the inhibitory effect of IAA. TZR promoted the growth of bulbils in *Lilium lancifolium* and *Dioscorea opposita* (Fan et al., 2019; Long et al., 1991), and IPA was important for the initiation of *Lilium lancifolium* bulbils (He et al., 2020). OPDA is the main precursor of jasmonic acid and might inhibit the initiation of bulbils in *Lilium lancifolium* and *Pinellia ternata* (Fan et al., 1999). In this study, the OPDA content was the lowest in the G2 stage and the highest in the G3 stage. The increase in OPDA content during the bulbils expansion stage might promote the maturation of bulbils. In this experiment, the soluble protein content rapidly increased in the G2 stage because it was the peak period of bulbils differentiation and required a large amount of protein. In the G3 and G4 stages, the bulbils continued to divide and the protein content remained high, which was similar to the research of Liu Wen(Liu, 2014). Currently, the transcriptome sequencing of *A. muelleri* is still in its infancy. Zheng et al. (2013) verified 177 polymorphic markers. Diao et al. (2014) obtained 80,332 differential genes. Zhong et al. (2018) obtained the neutral ceramidase in the sphingolipid metabolism pathway. The genes of *A. muelleri* foliar bulbils mostly increased in the early stage and decreased in the later stage. KEGG pathway enrichment analysis showed that the differential genes were concentrated in five metabolic pathways, among which the differences in starch and sucrose metabolism were the most significant, indicating that material and energy metabolism was crucial for bulbils formation.

## Author contributions

All authors contributed to the study conception and design. YX Liu and ZM Li was responsible for the collection of literature and manuscript writing. JD He, SY Ge and RJ Wang cultivated and prepared the plant materials. YQ Xie and WH Li measured the data. XW Wu revised the manuscript, supervision, and funding acquisition. All authors read and approved the final manuscript.

## Data Availability

The datasets generated during and/or analysed during the current study are available from the corresponding author on reasonable request.

## Acknowledgements

We would like to thank Dr. Mohamed A. A. Ahmed who is in Alexandria University for his assistance with English language and grammatical editing of the manuscript.

## Reference

Behera SS, Ray RC. 2017. Nutritional and potential health benefits of konjacglucomannan, a promising polysaccharide of elephant foot yam, *Amorphophallus konjac* K. Koch: A review. Food Reviews International 33(1), 22–43.

Borzenkova RA, Sobyanina EA, Pozdeeva AA, Yashkov M. 1998. Effect of phytohormones on starch synthesizing capacity in growing potato tubers. Russian Journal Of Plant Physiology 45(4), 472–480.

Cai YG, Duan L, Qin JF, Chen GA, Guo BL, Yang LH. 2020. Preliminary study on winter sowing and plastic film of feld for konjac cuttings. Journal of Guangxi Agriculture 35(5), 32–37.

Chang L, Xu YM, Xue JP. 2007. Formation of microtubers from the petiole of *Pinellia ternata*(Thunb.) Berit. *in vitro* and change of endogenous hormones content. Journal of Huazhong Agricultural University 26(5), 612–615.

Chen RJ. 2018. Study on the correlation between rhizome expansion and carbon and nitrogen metabobolism of wild *Poa*. L. Master’s thesis, Gansu Agricultural University.

Chen YB, Zhong GQ, Zhao QH, Teng JX. 2005. Effect of external secretion on forming multi-leaves and multi-tubers of *Amorphophallus konjac* K. Koch. Amino acids and biological resources (1): 35–37.

Diao Y, Yang C, Yan M, Zheng X, Jin S, Wang Y, Hu Z, Zhang J. 2014. *De novo* transcriptome and small RNA analyses of two *Amorphophallus* species. Plos One 9(4): e95428.

Fan JP, Wang B, Yan FX, Liu JS, Chen MX. 2019. Morphological characteristics and physiological changes of bulbils development in *Lilium lancifolium*. Journal of Northeast Agricultural University 50(2), 18–27.

Friml J. 2003. Auxin transport-shaping the plant. Current Opinion in Plant Biology 6(1), 7–12.

Gomez-Roldan V, Fermas S, Brewer PB, Puech-Pagès V, Dun EA, Pillot JP, Letisse F, Matusova R, Danoun S, Portais JC. 2008. Strigolactone inhibition of shoot branching. Nature 455(7210), 189–194.

Gu XH. 2022. Research on the mechanism of far-red light regulating shootbranching through hormone signals in tomato. Master’s thesis, Zhejiang University.

Guo YL, Wu M, Li RQ, Cai ZX, Zhang HB. 2022. Thermostable physically crosslinked cryogel from carboxymethylated konjac glucomannan fabricated by freeze-thawing. Food hydrocolloids 122, 107103.

He G, Yang P, Tang Y, Cao Y, Qi X, Xu L, Ming J. 2020. Mechanism of exogenous cytokinins inducing bulbil formation in *Lilium lancifolium in vitro*. Plant Cell Reputation 39, 861–872.

Hetterscheid W, Ittenbach S. 1996. Everything you always wanted to know about *Amorphophallus*, but were afraid to stick your nose into. Aroideana 19, 7–131.

Huang Q, Jin W, Ye S, Hu Y, Wang YT, Xu W, Li J, Li B. 2016. Comparative studies of konjac flours extracted from *Amorphophallus guripingensis* and *Amorphophallus rivirei*: Based on chemical analysis and rheology. Food Hydrocolloids 57, 209–216.

Irwin DL, Aarssen LW. 1996. Effects of nutrient level on cost and benefit of apical dominance in *Epilobium ciliatum*. American Midland Naturalist 136(1), 14–28.

Jiang, XR. 2022. Mining candidate genes for the first effective branch number in *Brassica* napus. Master’s theisis, Guizhou University.

Kebrom TH. 2017. A growing stem inhibits bud outgrowth-the overlooked theory of apical dominance. Frontiers in Plant Science 8, 1874.

Kobayashi S. Tsujihata S, Hibi N, Tsukamoto Y. 2002. Preparation and rheological characterization of carboxymethyl konjac glucomannan. Food Hydrocolloids 16(4), 289–294.

Liu C, Yang L, Yang Z, Ji YH. 2019a. Complete chloroplast genome of the economically important crop, Amorphophallus konjac (Araceae). Mitochondrial DNA Part B 4(1), 1097–1098.

Liu EX, Yang CZ, Liu JD, Jin SR, Harijati N, Hu ZL, Diao Y, Zhao LL. 2019b. Comparative analysis of complete chloroplast genome sequences of four major *Amorphophallus* species. Scientific Reports 9(1), 809.

Liu LL, An CC, Ye XM, Yuan JL, Wang YP, Zhang F. 2022. Relationships Among Apical Dominance of Potato Tuber, the Number of Main Stem and Yield Components. Journal of Nuclear Agricultural Sciences 36(02), 329–340.

Liu PY, Chen JF. 1986. Studies on the morphological development and growth trend of tuber of elephant foot yam(*Amorphophallus rivirei* Durieu and *A. albus* P. Y. Liu et J. F. Chen). Acta Horticulturae Sinica (04), 263–270.

Liu W. 2014. Studies on the morpholohical anatomy and the physiological and biochemical changes during the formation of microtubers in *Diocorea opposite* cv. Tiegun. Master’s thesis, Henan Normal University.

Long WH, Guo HC, Xiao GL, Wang Q. 2011. Variation of Endogenous Hormone and Carbohydrate Contents in Growing Yam Bulbils. Acta Horticulturae Sinica 38(04), 753–760.

Luo R, D YS, Sun YY, Cao ZJ. 2014. Morphological Observation and Anatomical Study on Bulbil Development of *Pinellia Ternata*. Acta Botanica Boreali-Occidentalia Sinica 34(9), 1776–1781.

Ongaro V, Leyser O. 2008. Hormonal control of shoot branching. Journal of Experimental Botany 59(1), 67–74.

Planas-Riverola A, Gupta A, Betegón-Putze I, Bosch N, Ibañes M, Caño-Delgado AI. 2019. Brassinosteroid signaling in plant development and adaptation to stress. Development 146(5).

Podwyszynska M. 2006. Improvement of bulb formation in micropropagated tulips by treatment with NAA and paclobutrazol or ancymidol. Acta Horticulturae 725, 679–684.

Qin X. 2004. The Research of Effect on growth with cutting off the terminal bud of *Amophophallus konjac*. Agricultural Education Research 19(1), 65.

Qiu CF. 2008. The study of breeding and cultivation on ornamental sweet potato variety. Master’s thesis, Fujian Agriculture and Forestry University.

Roumeliotis E, Kloosterman B, Oortwijn M, Kohlen W, Bouwmeester HJ, Visser RG, Bachem CW. 2012. The effects of auxin and strigolactones on tuber initiation and stolon architecture in potato. Journal of Experimental Botany 63(12), 4539–4547.

Ruegger M, Dewey E, Hobbie L, Brown D, Bernasconi P, Turner J, Muday G, Estelle M. 1997. Reduced naphthylphthalamic acid binding in the *tir3* mutant of *Arabidopsis* is associated with a reduction in polar auxin transport and diverse morphological defects. Plant Cell 9(5), 745–757.

Salam B B, Malka S K, Zhu XB, Gong HL, Ziv C, Teper-Bamnolker P, Ori N, Jiang JM, Eshel D. 2017. Etiolated stem branching is a result of systemic signaling associated with sucrose level. Plant Physiology 175(2), 734–745.

Schneider A, Godin C, Boudon F, Demotes-Mainard S, Sark S, Bertheloot J. 2019. Light regulation of axillary bud outgrowth along plant axes: an overview of the roles of sugars and hormones. Frontiers in Plant Science 10, 1296.

Shi CS. 2020. Excavation of critical genes for axillary bud germinationof *Pinus massoniana*. Master’s thesis, Guizhou University.

Shi YM, Tao YW, Lu YJ, Fei XN. 1998. Morphology of starch and mannan granules in corms of *Amorphophallus conjac*. Guihaia (1): 76–79+101.

Smirnova NI, Mestechkina NM, Shcherbukhin VD. 2002. Localization of acetyl groups in macromolecules of glucomannan from *Eremurus zangezuricus* roots. Prikladnaia Biokhimiia i Mikrobiologiia 38(5), 467–469.

Su YJ, Zhang MZ, Chang CH, Li JH, Sun YY, Cai YD, Xiong W, Gu LP, Yang YJ. 2022. The effect of citric-acid treatment on the physicochemical and gel properties of konjac glucomannan from *Amorphophallus bulbifer*. International Journal of Biological Macromolecules 216, 95–104.

Wang QY, Yang M, Wei HY, et al. 2022. Identification of *Alternaria* species causing leaf spot of *Amorphophallus bulbifer* in Yunnan province. Plant Protection 48(4), 240–244.

Wang TX, Wang RJ, Zhang LN, Xie YQ, Wu XW, Zhang D, Zheng L. 2023. Analysis of SSR and SNP loci in corm transcriptome of *Amorphophallus bulbifer* at different dormancy stages. Molecular Plant Breeding 22(15), 5007–5013.

Wang YP. 2021. Isolation and funtional analaysis of reactive oxygen species synthetic-related genes *TaRBOHF1*s in *Triticum aestivum* L. Master’s thesis, Shandong Agricultural University.

Wei HY, Wei W, Yang M, et al. 2020. Identification of a soft rot disease pathogen of *Amorphophallus bulbifer* in Yunnan Province. Acta Phytopathologica Sinica 50(04), 381–386.

Xia YT, Wang C, Hao N, Wu T. 2022. Research progress on plant branchiness and its application in vegetable breeding. China Vegetables (01): 31–40.

Zhang C, Chen JD, Yang FQ. 2014. Konjac glucomannan, a promising polysaccharide for OCDDS. Carbohydrate Polymers 104, 175–181.

Zhang DH, Wang QP, Duan ZB, Mi KX. 2009. Mechanism of relay multi-seedling release *Amorphophallus bulbifer* and its application in southeast Asia. Resource Development and Marketing 25(8), 682–684+710.

Zhang HY, Sun X, Yu H, Sun Q, Zhou L, Zhang YH. 2023. Effects of gibberellin and cytokinin on the tillers bud morphology of *Cynodon dactylon*. Pratacultural Science 40(05), 1368–1377.

Zhang W, Rhim JW. 2022. Recent progress in konjac glucomannan-based active food packaging films and property enhancement strategies. Food Hydrocolloids 128: 107572.

Zhang Y, Yu BS, Chen Y. 1994. Anatomical Study on the generation of the buibil of the Chinese yam. Journal of China Agricultural University 20(4): 413–418.

Zheng R, Wu Y, Xia Y. 2012. Chlorocholine chloride and paclobutrazol treatments promote carbohydrate accumulation in bulbs of *Lilium* oriental hybrids‘Sorbonne’. Journa of Zhejiang University-SCIENCE B(Biomedicine & Biotechnology) 13(2), 136–144.

Zheng X, Pan C, Diao Y, You Y, Yang C, Hu Z. 2013. Development of microsatellite markers by transcriptome sequencing in two species of *Amorphophallus* (Araceae). BMC Genomics 14, 490.

Zhou FZ, Wu ZK, Tian ZX, Zhang DC, Diao Y, Hu ZL. 2010. A new multi-leaves resource of *Amorphophallus konjac* K.Coch. Yangtze River Vegetables 274(20), 16–17.

Zhu YY, Luo R, Chen HL, Liu D. 2018. Bulbil development of *Pinellia cordata*. Guihaia 38(2):225–232.

